# Variation in the oxidative burst in response to wounding and bacterial infection among native and invading genotypes of yellow starthistle *(Centaurea solstitialis)*

**DOI:** 10.1101/156083

**Authors:** Angela M. Kaczowka, Patricia Lu-Irving, David A Baltrus, Katrina M. Dlugosch

## Abstract

- *Premise of the study:* Invasive plants may leave enemies behind when they colonize a new habitat, allowing selection to favor increased investment in growth and/or reproduction over defensive traits. Previous studies have identified reduced diversity of potential bacterial pathogens and evolutionary increases in growth and reproduction in invading populations of yellow starthistle (*Centaurea solstitialis*). This study leverages a recently developed high-throughput assay of immune function to test for evidence of a trade-off between increased growth and defense against bacterial pathogens in yellow starthistle’s invasion of California (USA).
- *Methods:* Seven bacterial strains were cultured from infected leaf tissue in the native range. Healthy leaf tissue from five native European collections and six invading collections were exposed to these native bacterial strains. A standardized assay of peroxidase activity was used measure the oxidative burst immune response to pathogen recognition by the leaf. Immune responses were compared to plant growth within and between ranges to assess evidence for a trade-off.
- *Key Results:* Plant genotypes from the native range demonstrated a higher immune response to bacterial strains than did invading genotypes, consistent with a trade-off with plant growth across regions. The same trade-off was also apparent across genotypes from the native range, but not across genotypes from the invaded range.
- *Conclusions:* Our results provide evidence that increased growth in a highly invasive plant species may come at a cost to immune function, consistent with the hypothesis that escape from enemies can provide opportunities for shifts in resource allocation that favor the proliferation of non-native species.

## INTRODUCTION

It is well documented that introduced plants can become invasive weeds that threaten the survival of native species, and cause significant environmental and economic damage (Pimentel et al., 2005; Pejchar and Mooney, 2009). Two related hypotheses are often invoked to explain why these exotic plants are so successful: the Enemy Release hypothesis and the Evolution of Increased Competitive Ability hypothesis. The Enemy Release hypothesis describes a scenario in which the invasive plant has left many of its natural enemies behind in the native range (Keane and Crawley, 2002; Mitchell et al., 2010). By escaping enemies, the invasive plant’s population is no longer regulated by its natural enemies, allowing explosive population growth in its new range. Often this hypothesis focuses on escape from herbivores, but escape from plant pathogens may also lead to increased success (Mitchell and Power, 2003; Colautti et al., 2004; Kulmatiski et al., 2008; Faillace et al., 2017). Furthermore, the Evolution of Increased Competitive Ability hypothesis suggests that escaping from enemies can facilitate adaptation, as selection may favor individuals that have increased their growth rate and reproductive output and in doing so sacrificed defensive mechanisms for enemies they no longer experience (Blossey and Notzold, 1995; Maron et al., 2004; Blumenthal et al., 2009; Gilbert and Parker, 2010). Growth-defense trade-offs are known to affect plants in general (Huot et al., 2014) and could potentially play a role in the success of invasive plants.

In this paper, we ask whether there is evidence of a growth-defense trade-off in the highly invasive weed, yellow starthistle (*Centaurea solstitialis)*. Specifically, we look for differences in plant pathogen immune responses between its native European range and its severe invasion of western North America. Yellow starthistle is native to Eurasia and was accidentally introduced to South America in the 1600’s and then to North America in the 1800’s in contaminated alfalfa seed (Gerlach, 1997). The invasions of both California (USA) and South America are derived from western European genotypes in the native range (Fig. 1A; (Barker et al., 2017)). Yellow starthistle is a colonizer of grassland systems and has invaded over 14 million acres in California (Pitcairn et al., 2006). The state of California has been spending approximately $12.5 million each year on controlling this invasion (Ditomaso et al., 2006; Pitcairn et al., 2006).

**Figure 1.**
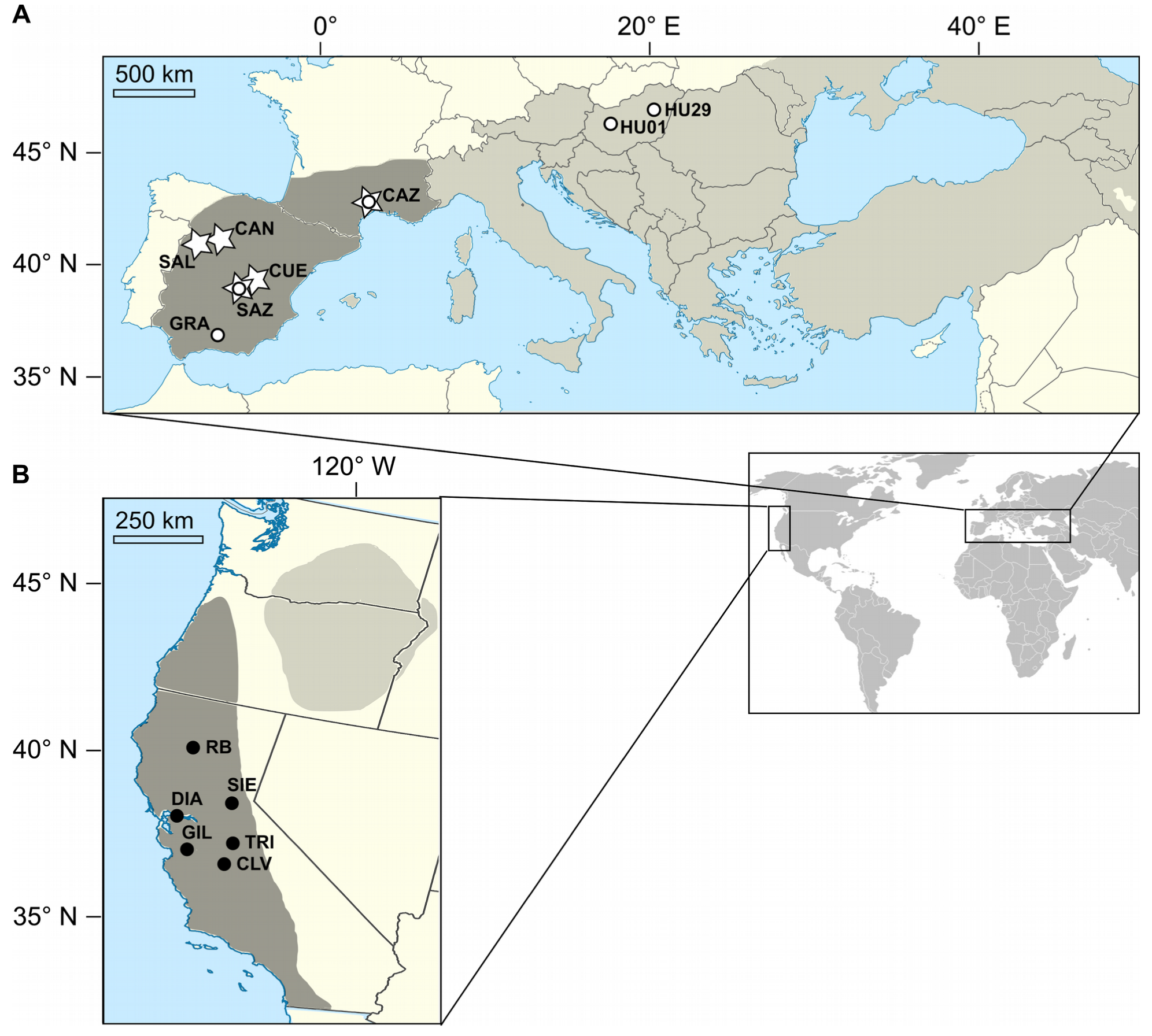
The distribution (gray) of yellow starthistle and sampling sites for this study. Maps detail the native range in Eurasia (a) and the invasion of western North America (b). Previous work has indicated that western Europe is the source for the invasion of California, USA (both in dark shading; Barker et al. 2017). Sampling for plant genotypes included six locations in California (B, filled circles), three locations in western Europe and an additional two locations in eastern Europe (A, open circles). Sampling for bacterial genotypes included five sites in western Europe (A, stars).

Previous work on yellow starthistle has demonstrated that there are significant differences between the microbiome of yellow starthistle in the native range and the microbiome of plants in the invaded range. Bacterial communities in the microbiome of the California invaders had significantly lower diversity than in the native plants (Lu-Irving et al., 2017), and plants have more favorable interactions with the soil microbial community in the California invasion when compared to soil communities from other ranges (Andonian and Hierro, 2011; Andonian et al., 2011, 2012). There are also life history differences between the plants in the native and invaded ranges. When genotypes from both the native and invaded range are grown in a common environment, the invasive genotypes grow larger above and belowground (Widmer et al., 2007; Eriksen et al., 2012; Dlugosch et al., 2015). The invasive genotypes also have earlier flowering times and increased reproductive output (Dlugosch et al., 2015). Taken together, this information suggests that increased growth could be a benefit of escape from bacterial pathogens in this system.

Plant immune response consists of two different systems, one that occurs between the membranes (transmembrane pattern recognition receptors, PRRs) and one that occurs within the cell using NB-LRR protein products as a result of specific resistance ‘R’ genes (Jones and Dangl, 2006). When plant pathogens are exposed to plant cells, they release nonprotein and protein effectors called pathogen associated molecular patterns (PAMPs) which are recognized by the PRRs (Medzhitov, 2013). This recognition elicits PAMP triggered immunity (PTI) in the plant (Chisholm et al., 2006). Pathogens can interfere with PTI by releasing effectors resulting in effector triggered susceptibility (ETS) (Abramovitch and Martin, 2004; Dodds and Rathjen, 2010; Win et al., 2012). If these effectors are directly or indirectly recognized by one of the NB-LRR proteins within the plant cell, effector triggered immunity (ETI) occurs within the plant (Chisholm et al., 2006; Jones and Dangl, 2006). This is a much stronger response than the PTI and is often called a hypersensitive response (HR), which typically results in cell death at the site of the infection (Chisholm et al., 2006; Jones and Dangl, 2006; Camejo et al., 2016). Both PTI and HR will alter hormone activity in the plant and can actively suppress growth (Huot et al., 2014).

Here we assay these immune responses in yellow starthistle by examining the ‘oxidative burst’ that is created during PTI and HR (Lamb and Dixon, 1997; Camejo et al., 2016). This burst is a localized production of reactive oxygen species (ROS) used to hinder pathogen invasion (Wojtaszek, 1997). Pathogen attacks elicit ROS production from NADPH oxidase apoplastic peroxidases (Wojtaszek, 1997; Chakravarthy et al., 2009; Daudi et al., 2012; Camejo et al., 2016). The ROS consists of **superoxide radical** (**·**O_2_^−^), hydrogen peroxide (H_2_O_2_) and hydroxyl radical (**·**OH) which are released from NADPH oxidase and the apoplastic peroxidase enzymes (Lamb and Dixon, 1997; Wojtaszek, 1997; Camejo et al., 2016). The apoplastic peroxidase enzymes have previously been used to measure plant immune response to pathogens, using an assay of peroxidase activity (Mott et al., 2016). This assay measures peroxidase activity in response to plant pathogen sensitivity through a visible color change. Substrates added during the assay will produce a brown color when acted on by peroxidase which can be quantified by reading absorbance at 550 nm (Mott et al., 2016). By using this technique to assay plant responses to native bacterial pathogens, we ask whether there is a loss of immune activity in the invading genotypes. A lower immune response in these high growth genotypes would support the possibility that the fitness of invasive plants has been increased as a result of a growth-defense trade-off.

## MATERIALS AND METHODS

### Bacterial collections

Bacterial strains were obtained from field-collected leaf material in the native range. Living leaves on the basal rosette with evidence of disease symptoms were collected in July 2015 from five field sites in western Europe (Fig. 1; Table 1) and stored in sterile falcon tubes. The remainder of the plant was pressed and dried after sampling, and submitted to the University of Arizona Herbarium (ARIZ; Table 2). Samples were kept on ice in the field and stored at 4 °C. Leaves were removed from tubes using sterile forceps. Forceps were sterilized in a 200ppm bleach for 2 minutes, and then rinsed with sterile water. To obtain bacterial strains in culture, a 1 cm^2^ piece of leaf was cut out using a sterile scalpel. This leaf tissue was placed in a 1.5mL tube containing 400 μL sterile 10mM MgCl_2_ and macerated for 45 seconds using a sterile plastic pellet pestle. An additional 600 μL sterile 10mM MgCl_2_ was added to bring the volume of each tube to 1mL, and 150μL of macerated leaf suspension was spread on malt extract agar plate using a sterile plastic cell spreader. Two 10-fold serial dilutions were made and 10 μL of each dilution was spotted on a second malt agar plate.

**Table 1.** Locality information for seed and bacterial collections.

| Site code | Nearest city | Country/State | Continent | Lat. | Long. | Seed Source | Strain source |
| --- | --- | --- | --- | --- | --- | --- | --- |
| RB | Red Bluff | Northern California | North America | 40.271 | −122.271 | X |  |
| SIE | Placerville | California Sierra Nevadas | North America | 38.782 | −120.416 | X |  |
| DIA | Clayton | California coast | North America | 37.865 | −121.978 | X |  |
| TRI | Mariposa | California Sierra Nevadas | North America | 37.462 | −119.792 | X |  |
| GIL | Gilroy | California coast | North America | 37.034 | −121.537 | X |  |
| CLV | Clovis | California valley | North America | 36.916 | −119.794 | X |  |
| SAL | Salamanca | Spain | Eurasia | 40.99 | −5.659 |  | X |
| CAN | Canales | Spain | Eurasia | 41 | −4.897 |  | X |
| GRA | Granada | Spain | Eurasia | 37.268 | −3.665 | X |  |
| SAZ | Saz | Spain | Eurasia | 39.835 | −2.511 |  | X |
| CUE | Cuenca | Spain | Eurasia | 40.129 | −2.140 |  | X |
| CAZ | Caze | France | Eurasia | 43.748 | 3.771 | X | X |
| HU01 | Várpalota | Hungary | Eurasia | 47.182 | 18.089 | X |  |
| HU29 | Püspökládany | Hungary | Eurasia | 47.318 | 21.034 | X |  |

**Table 2.** Source location, taxonomic identification, and herbarium (ARIZ) accession for each bacterial strain.

| Field Site | Strain Code | Taxonomic name | Accession Number |
| --- | --- | --- | --- |
| CAN | 1031 | *Pseudomonas viridiflava* | CAN201507 |
| CAZ | 1032 | *Pseudomonas sp.* | CAZ201509 |
| SAL | 1033 | *Pseudomonas asturiensis* | SAL201503 |
| SAZ | 1038 | *Curtobacterium flaccumfaciens* | SAZ201521 |
| CUE | 1042 | *Massilia sp.* | CUE201501 |
| CUE | 1043 | *Curtobacterium flaccumfaciens* | CUE201501 |
| CAN | 1044 | *Pseudomonas congelans* | CAN201511 |

Plates were incubated on lab bench under low light at approximately ~27 °C for 2 days to allow bacterial cultures to grow. Single bacterial colonies were selected from the plates and diluted in 25 μL of sterile H_2_O for DNA identification and storage. For storage, single colony dilutions were cultured in 2mL LB media, and 750 μL of liquid culture was mixed with 750 μL of autoclaved 80% glycerol and stored at −80 °C.

Bacteria were identified by sequencing the 16S rRNA region of DNA, using universal primers 27F (5’TACGGYTACCTTGTTACGACTT3’) and 149R (5’AGAGTTTGATCMTGGCTCAG3’). Amplification was performed in a 50uL reaction containing 1 μL of bacterial single colony dilution, 2.5 μL of each primer at 10 μm, 0.625 μL Taq, 1.25 μL of 5mM dNTPs, and 50μL of 10x PCR buffer. Reaction conditions included 95 °C for 5 minutes, and 35 cycles of 94 °C for 15 seconds, 50 °C for 30 seconds, and 72 °C for 1 minute 30 seconds. PCR products were Sanger sequenced by Genoscreen (Lille, France), and sequences were compared to existing sequences on GenBank for taxonomic identification.

### Plant genotype collections

Plant genotypes were collected in 2008 from six sites in the invaded range and five sites in the native range (Fig. 1; Table 1). Distinct genetic subpopulations have been identified in the native range, including those in Mediterranean western Europe, central-eastern Europe, Asia (including the Middle East), and the Balkan-Apennine peninsulas (Barker et al., 2017). The invasion of California as well as invasions of South America appear to be derived almost entirely from western European genotypes (Fig. 1; Barker et al. 2017), and our sampling here includes genotypes from both the western European and the central-eastern European subpopulations. At each site, seeds were collected from multiple maternal plants separated by at least one meter along a linear transect.

Seeds from at least 5 maternal plants per population were germinated in flats filled with moistened soil [3:2:1 ratio of Sunshine Mix #3 soil (SunGro Horticulture), vermiculite, and 20 grit silica sand]. Plant Preservative (1% v/v; Plant Cell Technology) was sprayed on top of seeds to minimize fungus growth. Seedlings were reared for four weeks under artificial light in laboratory conditions, and then transplanted into 410 ml Deepots (Stuewe & Sons) and moved to glasshouse conditions at the University of Arizona under a 14 hour daylength.

### Peroxidase assay

Petri dishes were filled with LB agar and streaked with frozen bacterial cultures to allow isolation of single bacterial colonies from the stored cultures. Plates were placed in a 27° C incubator for approximately 36 hours. Single bacterial colonies were selected and used to inoculate 3mL liquid LB cultures of each strain. Test tubes containing the live liquid cultures were then placed in a 27 °C shaker for approximately 44 hours. Live liquid cultures were then diluted by mixing 25μL of live strain with 25μL of LB in a microfuge tube. Microfuge tubes containing diluted strains were thoroughly mixed before use in the peroxidase assay.

Healthy, basal leaves were removed from each individual plant and wiped off with a paper towel sprayed with 70% ethanol. Leaf discs were obtained with a 4.8 mm diameter cork borer, avoiding major veins (Fig. 3A). Each leaf was photographed after sampling. Leaf discs were placed inside a 1.5mL microfuge tube and filled with 1mL of 1X Murashige & Skoog basal salt mixture (MS) buffer. The microfuge tubes were shaken in a 27° C shaker for one hour.

We used a modified version of the peroxidase assay protocol by Mott and colleagues (Mott et al., 2016). The inner 60 wells of flat bottom 96 well microliter plates were filled with 50μl of 1 X MS. The outer edges of the wells were filled with 300μl of ddH2O (Millipore water) to minimize evaporation and edge effects. Leaf discs were placed in the bottom of each well, following a diagonal plate design to prevent spatial effects (Fig 2; Fig 3B). Each combination of plant genotype and bacterial strain was replicated 2-4 times, and additional wells were used for controls including ‘No Leaf’ (bacteria but no leaf disk), ‘Leaf Only’ (leaf disk with no bacteria), and ‘Media Only’ (neither leaf disk nor bacteria). Each well including a strain was treated with diluted 1μL of culture. The 96 well-plates were covered with breathable sealing film. Sealed plates were stacked and stored in containers lined with wet paper towels to limit evaporation. Containers were placed in a 27°C shaker for 16 hours. All leaf discs were removed from the plates; forceps were rinsed with Millipore water in between each well. After all the leaf discs were removed, plates were placed into a Synergy H1 Hybrid Multi-Mode Microplate Reader (BioTek) to record any initial absorbance of the solution. 50μl of 1 mg/mL solution of 5-aminosalicylic acid (5-AS) with 0.01% hydrogen peroxide was added to each well. After one minute, 20μl of 2N NaOH was added to each well to stop the color changing reaction (Fig 3C). Plates were placed into the H1 Hybrid Reader to record the final absorbance of each well. Absorbance values for the second experiment are differences between final readings and initial readings.

**Figure 2.**
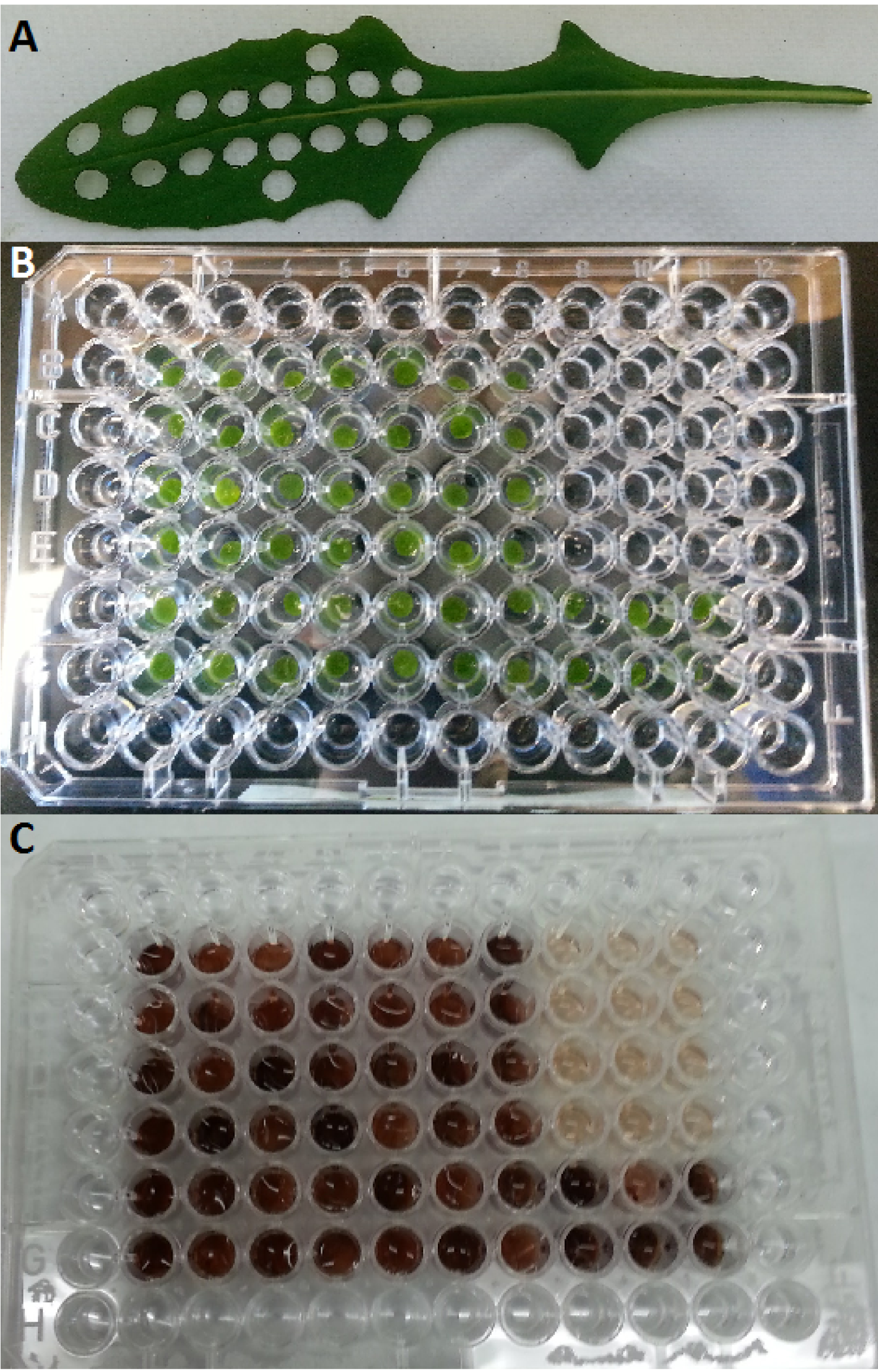
A) Leaf after sampling. B) Plate with incubated leaf disks, bacterial strains, and controls. C) Plate after peroxidase assay complete.

**Figure 3.**
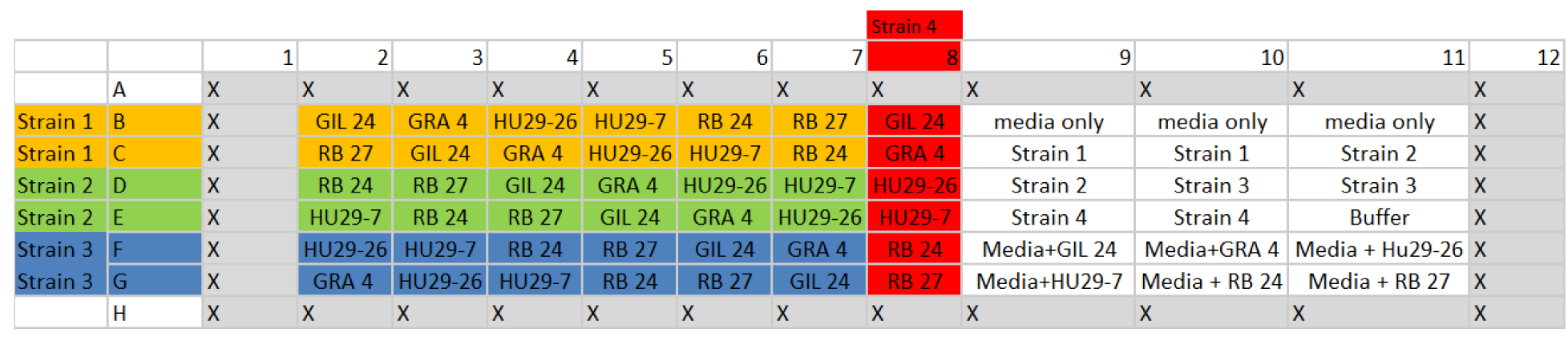
Example of plate design. Each plate included four strains combined with 12 individuals from at least four populations, as well as separate controls for each strain and individual.

### Experiment 1: Method development

An initial experiment was performed to optimize assay conditions. Three plant genotypes were selected for use in this experiment, GRA (n=3), GIL (n=3), and CAZ (n=1), and exposed to strains 1031 (*Pseudomonas viridiflava*) and 1042 (*Massilia sp*). The peroxidase assay was repeated with autoclaved and live samples of strains 1031 and 1042, to determine whether autoclaved cultures were sufficient to elicit plant immune responses.

We evaluated whether plant immune response differed between live strains and autoclaved strains using a linear model with fixed effects of treatment, plant population nested within treatment, and plant individual nested within population and treatment. This nested design was necessary because plant populations were not present in the media only control treatment. Fixed effects of plant population and individual were used in correlations with fitness variables described below. We used Tukey’s honest significant difference (HSD) tests to identify differences in immune response between the live strain (1031, 1042) treatment, autoclaved strain (1031, 1042) treatment, ‘Leaf Only’ control treatment (containing a leaf disk and sterile media for all genotypes), and the ‘No Leaf’ treatment (containing either sterile media, or bacterial strain in media, or only MS Buffer).

All statistical analyses were performed in JMP 13.1.0 (SAS Institute 2016).

### Experiment 2: Variation in immune response among plant genotypes

Eleven plant genotypes were selected to compare immune responses between native and invading regions: CAZ (n=7), CLV (n=7), DIA (n=6), GIL (n=8), GRA (n=3), HU01 (n=3), HU29 (n=7), RB (n=8), SAZ (n=3), SIE (n=2), TRI (n=6). These plants were exposed to the following strains during the assay: 1031 (*Pseudomonas viridiflava*), 1032 (*Pseudomonas sp*.), 1033 (*Pseudomonas asturiensis*), 1038 (*Curtobacterium flaccumfaciens*), 1042 (*Massilia sp*), 1043 (*Curtobacterium flaccumfaciens*), 1044 (*Pseudomonas congelans*). We evaluated whether yellow starthistle immune response differed amongst the seven strains and our controls using a linear model with fixed effects of treatment, population nested within treatment, and individual plant nested within treatment and population, and in Experiment 1. We used Tukey’s (HSD) tests to identify significant differences in immune response across treatments.

We tested for overall differences in immune response between the native and invasive plant genotypes. We used a linear model predicting absorbance with fixed effects of strain and region (native vs. invaded), and their interaction and nested effects of population within region, and individual plant within population and region. Fixed effects of plant population and individual were used in correlations with fitness variables described below. To evaluate how different yellow starthistle genotypes varied in their immune response to the seven bacterial strains (i.e to allow inspection of strain x plant population interactions), linear models were used to predict absorbance for each of the seven strains individually, with fixed effects of region, population nested within region, and individual nested within population and region.Tukey’s HSD tests were used to identify significant differences in immune response among populations.

### Trade-offs between immune response and plant size

Population means for peroxidase activity were compared to previously published data from a large common environment experiment, which identified evolutionary increases in growth and reproductive potential in the Californian invasion, and included the seed collections using in the present study (Dlugosch et al., 2015). Briefly, single offspring from 7–20 different mothers per site were reared in a randomized glasshouse common garden, and their size at 5.5 weeks measured using a linear morphological index that strongly correlates with biomass (Dlugosch et al., 2015): Size Index = [leaf number * (maximum leaf length*maximum leaf width)^1/2^]. Half of the plants per population were maintained through senescence and their date of first flower and total flowering head production recorded. The date of first flowering strongly predicted total reproductive output, such that invading plants flowered earlier, resulting in greater production of flowering heads (Dlugosch et al., 2015).

Population means for plant size from the previous study were used to predict mean population absorbance (Least Squares Means from analyses of population variation in absorbance detailed above) using linear models with fixed effects of strain, trait, and their interaction. Only strains showing significant differences in peroxidase activity between native and invaded regions (i.e. the potential for a trade-off with fitness) were used for these analyses. Given that regional differences in plant traits provide only a single comparison (native vs. invaded range) for evaluating potential trade-offs, models were fit within native and invaded regions separately to test for evidence of consistent trade-offs between plant size and immune function at this scale. Future analyses are planned at the population and individual level for the plants used in this study, when they complete flowering and are harvested for biomass measurements in the Summer of 2017.

## RESULTS

### Bacterial collections

Seven bacterial strains from field collections in the native range were utilized in this study (Table 2). These included four *Pseudomonas* taxa, two unique strains of *Curtobacterium flaccumfaciens*, and an unidentified species of *Massilia*.

### Experiment 1: Method development

This experiment demonstrated the importance of using live strains in our assay of plant immune response (Fig. 4A). Absorbance differed significantly among treatments (F_5,5_ = 7.3, *P* < 0.0001). Control treatments including no leaf disks (buffer, media, and strain only wells) showed the lowest peroxidase activity, followed by the ‘Leaf Only’ control treatment, and these two controls were not significantly different. Both autoclaved strain treatments and the live 1042 treatment had similar mean immune response (absorbance) as the ‘Leaf Only’ control. Only the live 1031 treatment was significantly different from both controls, showing a higher mean immune response than both the autoclaved treatments and the live 1042 treatment (Fig. 4A).

**Figure 4.**
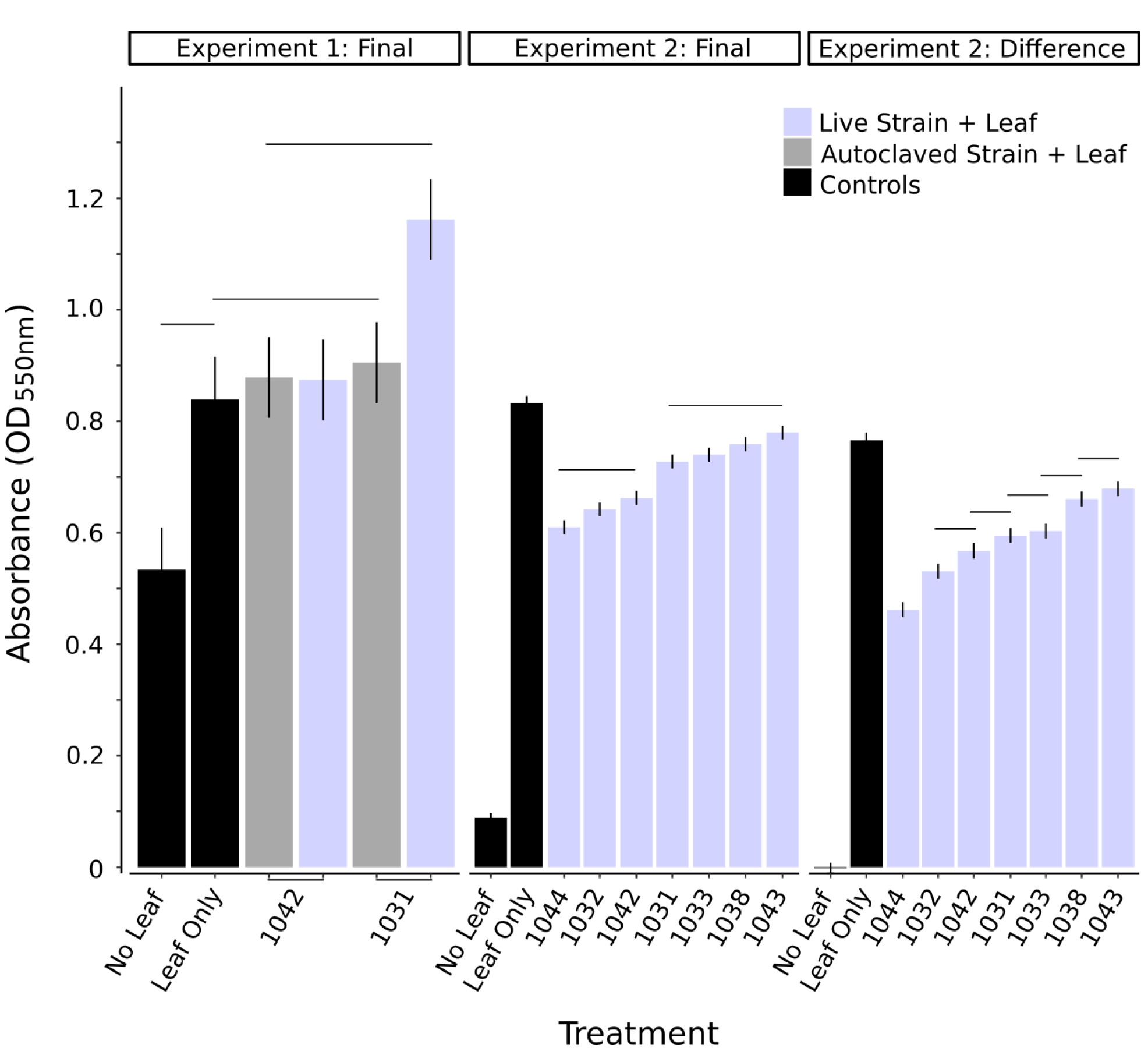
Control and strain treatments across all plant genotypes. Shown are Least Squares Means +/- s.e.m. from linear models predicting absorbance as a function of control and strain treatments. Horizontal lines connect treatments that were not significantly different. In Experiment 1, leaf disks were incubated with both autoclaved and live cultures of two bacterial strains. In Experiment 2, leaf disks were incubated with seven live strains. Control treatments included ‘No Leaf’ (containing either sterile media alone or a bacterial strain in media or MS buffer) and ‘Leaf Only’ (containing a leaf disk + sterile media for all leaf genotypes).

### Experiment 2: Variation in immune response among plant genotypes

In this experiment, initial absorbance readings were taken before the color change assay, and subtracting these values reduced variability within treatments from final readings alone (Fig. 4B,C). In particular, ‘No Leaf’ controls showed almost no evidence of absorbance after initial readings were removed (Fig. 4C). Treatment effects were significant overall (F_8,701_ = 457.5, *P* < 0.001) and indicated that all strains were different from controls (Fig. 4C). In contrast to results from the first experiment, the highest immune response was exhibited by the ‘Leaf Only’ control in this experiment, which was significantly higher than all other treatments. A separate analysis of the ‘Leaf Only’ treatment revealed no significant difference in absorbance between regions (F_1,60_ = 2.04, *P* = 0.16). Population effects were significant (F_9,49_ = 2.6, *P* = 0.013), but only indicated differences in Leaf Only absorbance between the most extreme populations across the regions (Fig. 5). While the high peroxidase activity in this treatment was unexpected and its source is not known (see Discussion), the lack of regional differences should indicate that it does not confound analyses of plant responses to bacterial strains.

**Figure 5.**
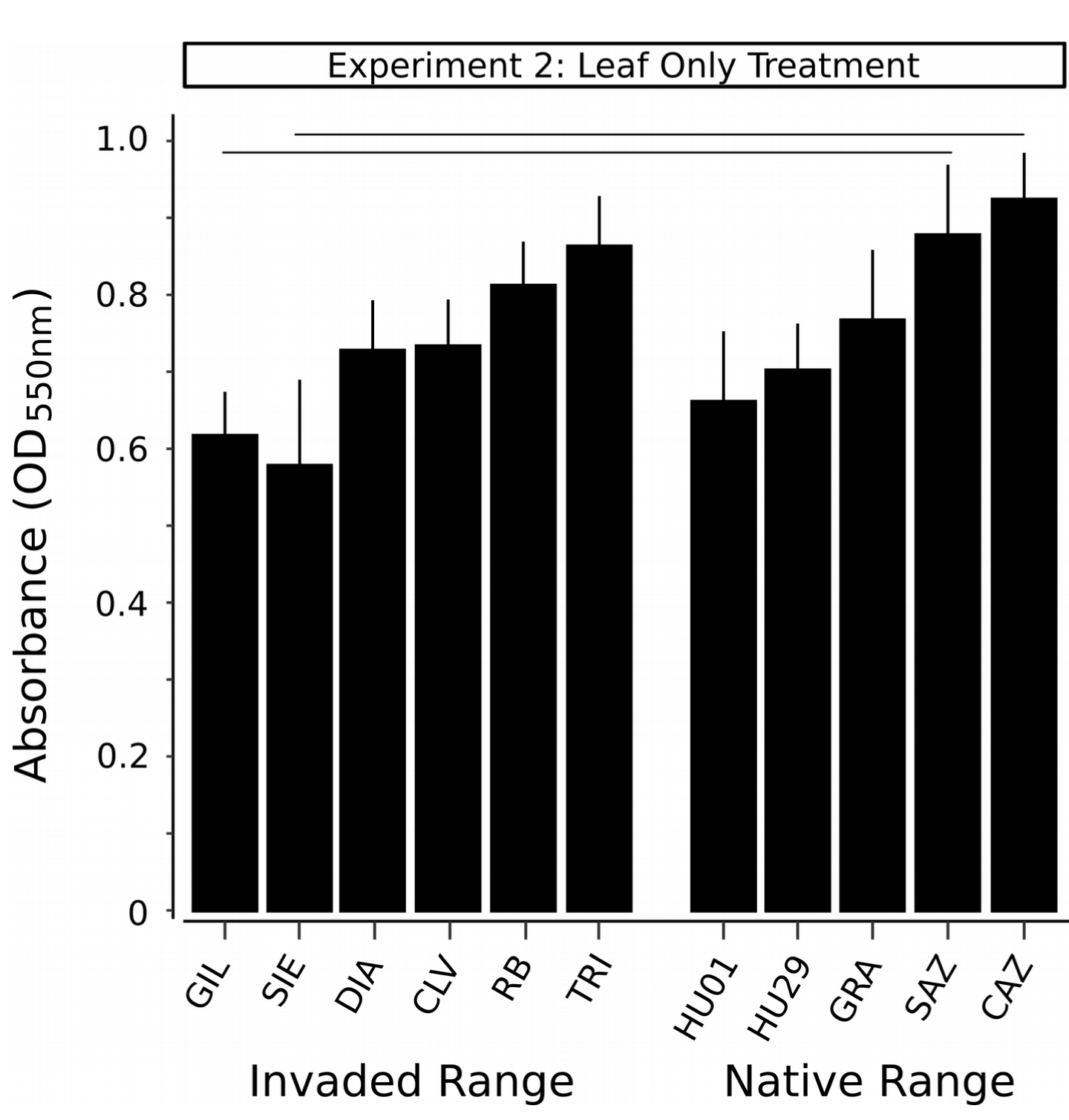
Leaf Only control treatment (containing a leaf disk + sterile media) for each plant source population. Shown are Least Squares Means +/- s.e.m. from a linear model predicting absorbance as a function of plant region (Invaded vs. Native; not significant, *P* = 0.16) and population (*P* = 0.01). Populations are ordered according to means and significant differences, and horizontal lines connect populations that are not significantly different. Absorbance values are differences between final readings and initial readings before the addition of 5-AS to assay the activity of peroxidase.

When leaf disks were combined with live bacterial strains, significant regional differences became apparent (Fig. 6). Absorbance was significantly affected by region (F_1,762_ = 13.2, *P* = 0.0001), strain (F_6,762_ = 16.6, *P* < 0.0001), and their interaction (Fig. 7; F_6,762_ = 2.3, *P* = 0.035), as well as nested effects of population within region (F_9,762_ = 12.5, *P* < 0.0001) and individual plant genotype within population (F_49,762_ = 7.9, *P* < 0.0001). Native plant genotypes had a higher mean immune response to all strains other than 1032 and 1043 (Fig. 6), and these latter two strains showed no significant differences between regions (Fig. 8). When exposed to a strain there were differences within region among genotypes (Fig. 7). Genotypes GIL and SIE (invaded range) and HU29 (native range) had the lowest immune response whereas RB (invaded range) and GRA and CAZ (native range) had the highest immune responses. Overall, the native genotype GRA had the highest immune response to the seven strains (Fig. 7).

**Figure 6.**
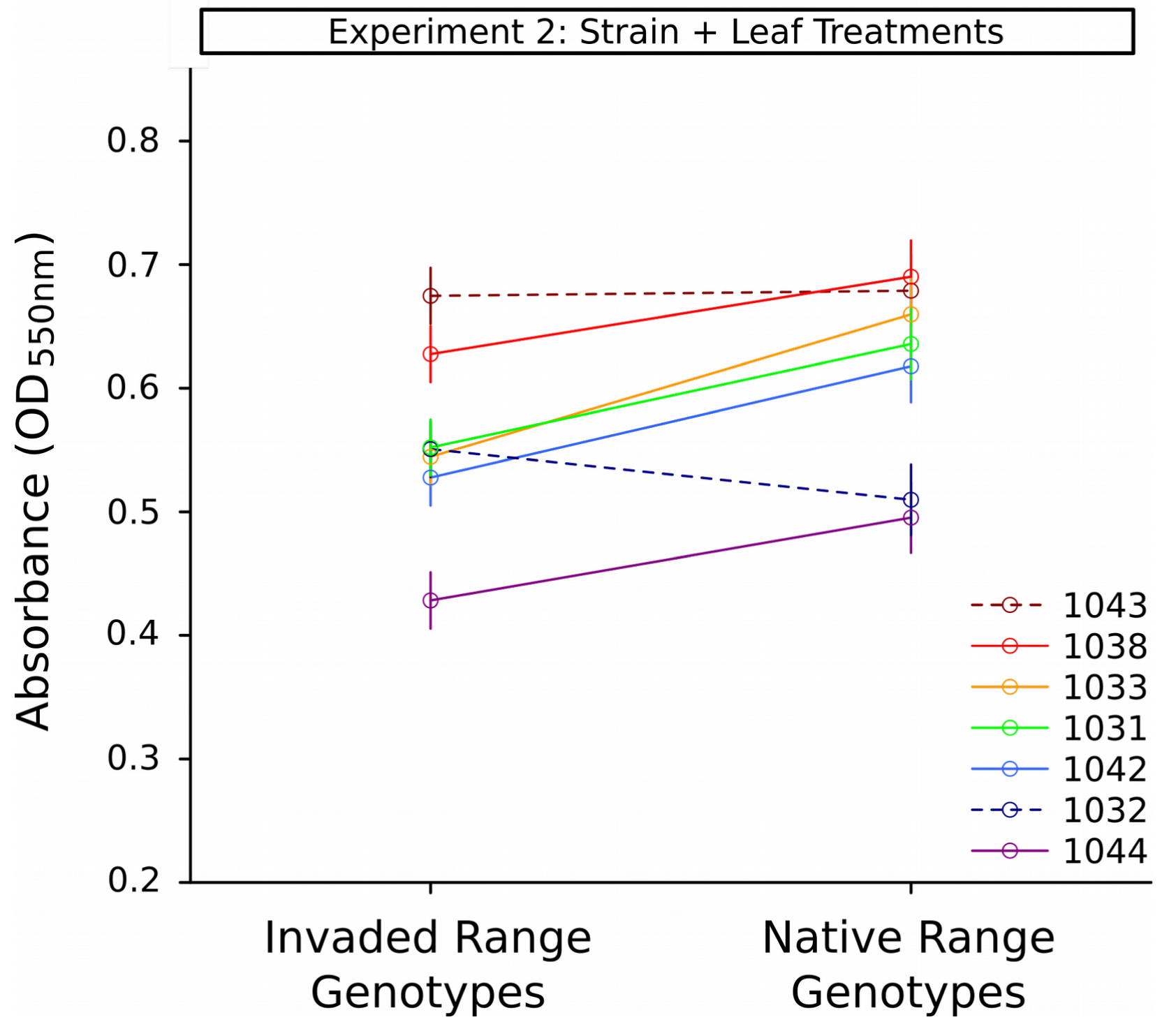
Interaction between the effect of plant genotype source region (Invaded vs. Native) and bacterial strain on absorbance in the peroxidase activity assay. Shown are Least Squares Means +/- s.e.m. from the linear model interaction term effects (*P* = 0.04). Bacterial strains are indicated by colors. Strains that showed significant or marginally significant differences between regions are indicated with solid lines, and those with no significant difference between regions are indicated with dashed lines. Absorbance values are differences between final readings and initial readings before the addition of 5-AS to assay the activity of peroxidase.

**Figure 7.**
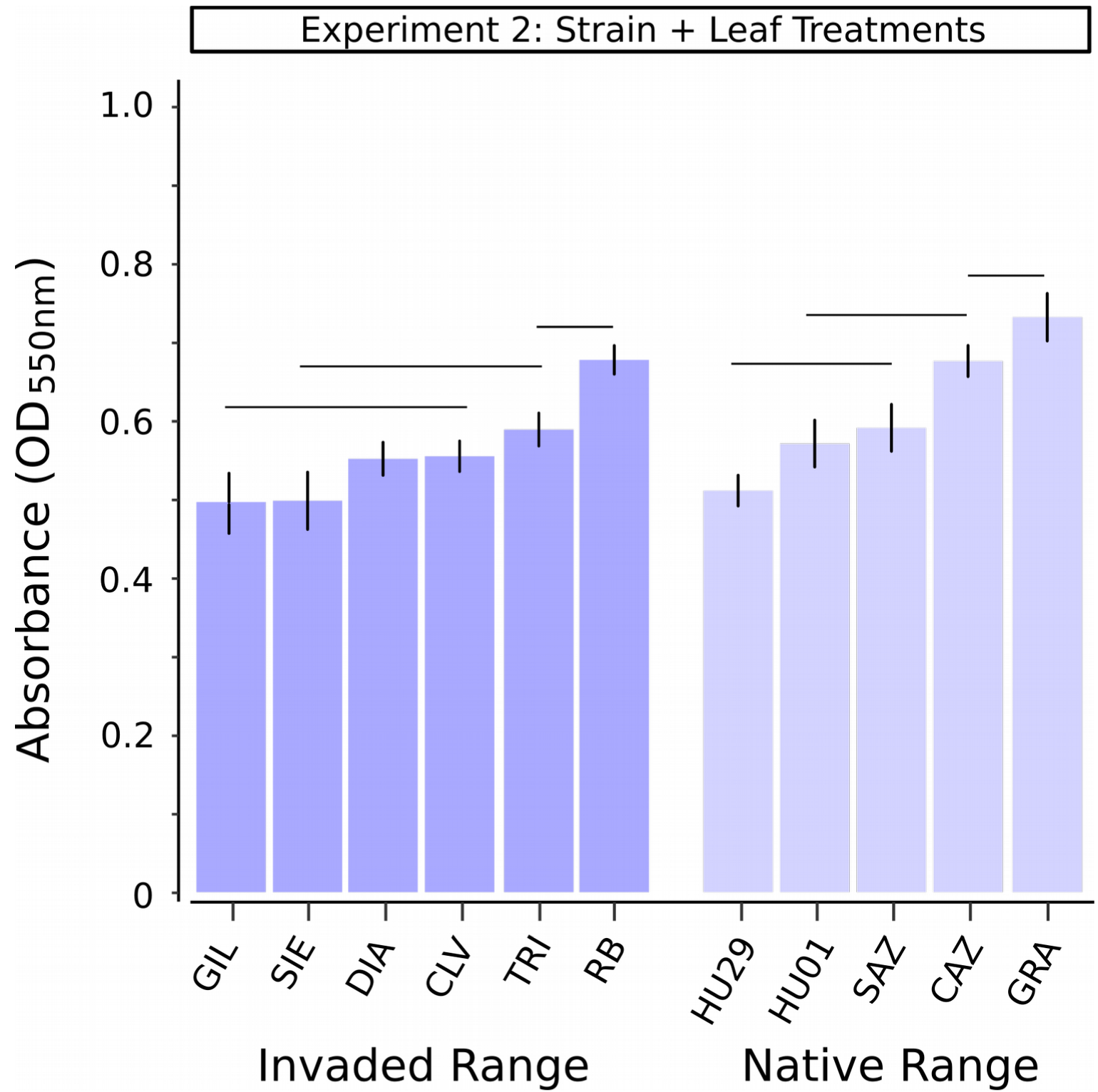
Plant source population effects across Strain+Leaf disk treatments. Shown are Least Squares Means +/- s.e.m. from a linear model predicting absorbance as a function of plant region (Invaded vs. Native; *P* = 0.0003), bacterial strain (*P* < 0.0001), region by strain interactions (*P* = 0.04), and populations within regions (*P* < 0.0001). Populations are ordered according to means and significant differences within regions, and horizontal lines connect populations that are not significantly different. Absorbance values are differences between final readings and initial readings before the addition of 5-AS to assay the activity of peroxidase.

**Figure 8.**
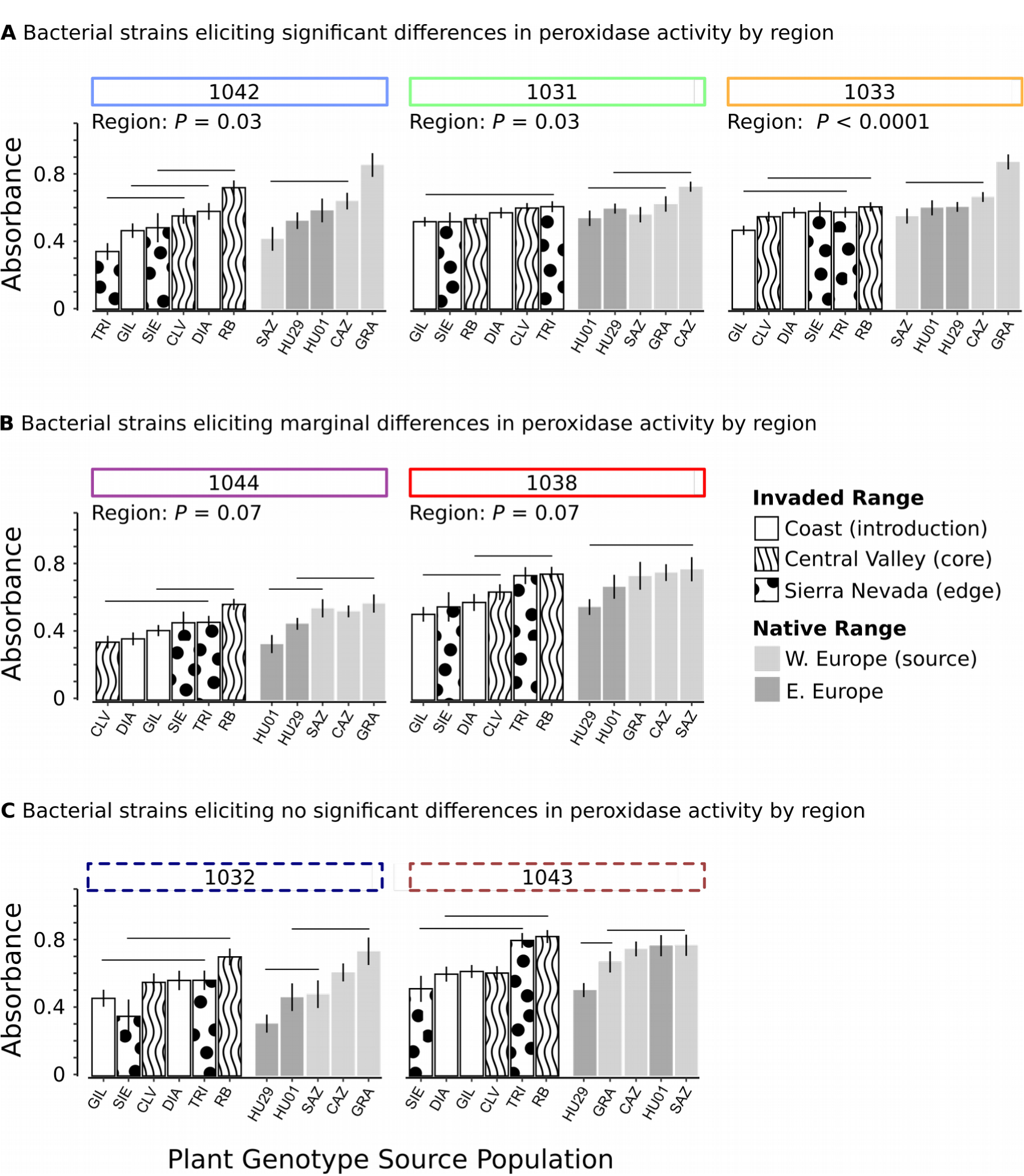
Plant source population effects in response to each strain in Strain+Leaf disk treatments. Shown are Least Squares Means +/- s.e.m. from a linear model predicting absorbance as a function of plant region and populations within regions. Populations are ordered according to means and significant differences within regions, and horizontal lines connect populations that are not significantly different. Absorbance values are differences between final readings and initial readings before the addition of 5-AS to assay the activity of peroxidase. All strains showed significant differences among populations within regions (*P* < 0.01 in each case), and two strains showed no significant differences between native and invaded region genotypes (C; 1032: *P* = 0.8; 1043: *P* = 0.3).

Population effects revealed that the highest immune responses within the native range were observed in populations from western Europe (Fig. 7), which is the source region for the invasion of California (Fig 1). Within the invasion, populations also differed significantly (Fig. 7), with the lowest immune responses generally found in coastal populations near the original introduction (DIA, GIL) and a population near the eastern leading edge of expansion in the Sierra Nevada mountains (SIE). The highest response was observed in a population from the northern Central Valley region (RB). Inspecting population variation within strains, these regional patterns were generally consistent and did not vary substantially by strain (Fig. 8).

### Trade-offs between immune response and plant size

All five strains that elicited significant or marginally significant regional differences showed lower mean immune response in genotypes from the invade range (Fig. 6), suggesting a potential negative trade-off between growth in the absence of pathogens and defense function across regions (Fig. 9B). Consistent with a trade-off, variation in peroxidase activity within the native range was significantly negatively related to plant size as measured in the previously published glasshouse experiment (Fig. 9C). Mean population absorbance in the native range was predicted by significant fixed effects of plant size (F_1,19_ = 12.1, *P* = 0.003) and strain (F_4,19_ = 3.8, *P* = 0.02). A non-significant interaction of population and strain (*P* = 0.9) was removed from the final model. In contrast, within the invaded range there was no significant relationship between peroxidase activity and plant size (Fig 9D; Size F_1,24_ = 0.77, *P* = 0.4; Strain F_4,24_ = 4.0, *P* = 0.01), with population mean absorbance values across the invaded range generally similar to the values demonstrated by the native populations with the largest size phenotypes, for a given bacterial strain (Fig. 9C,D).

**Figure 9.**
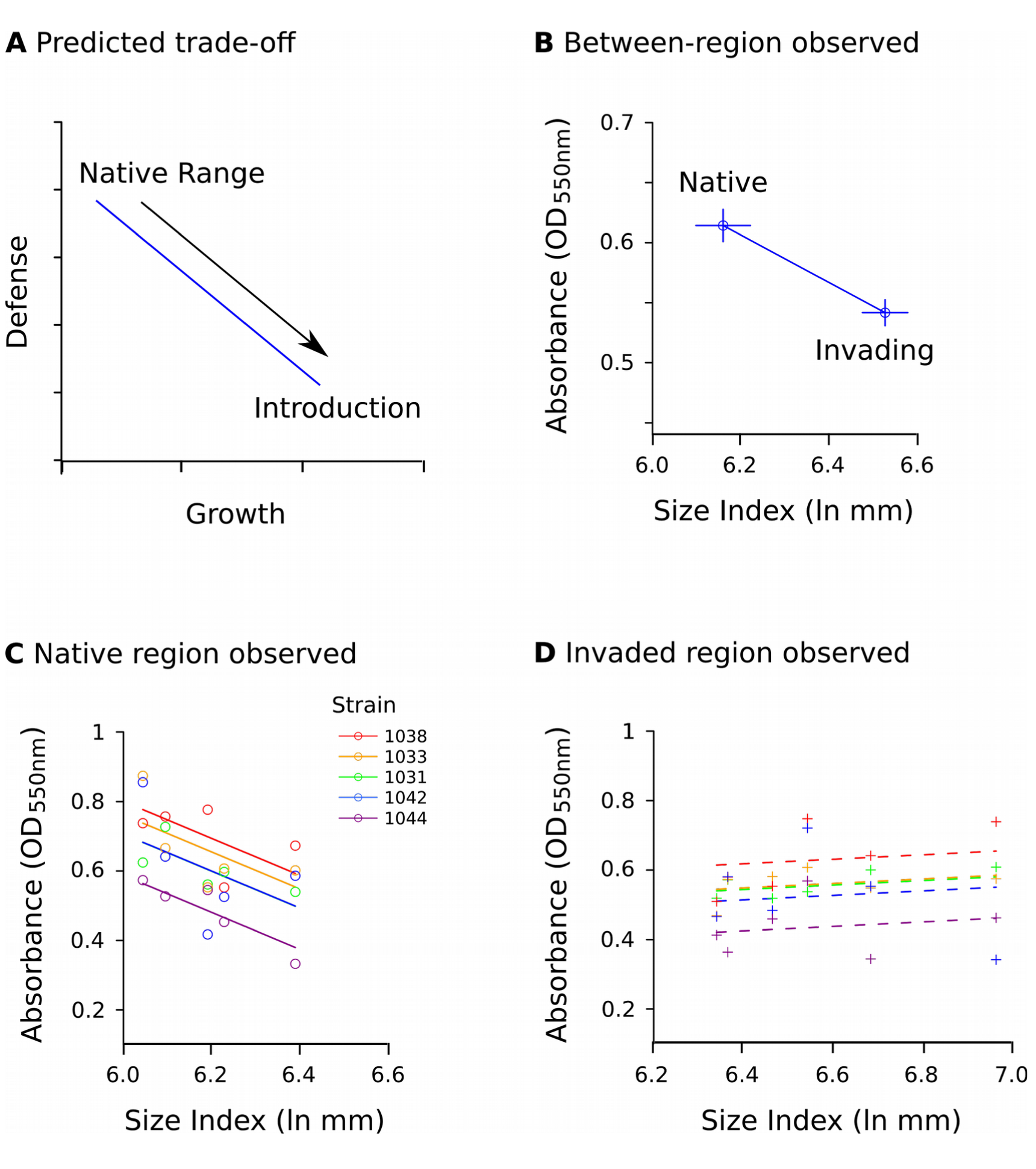
A) Predicted trade-off between investment in defense functions and growth in the absence of enemies. If introduced populations escape enemies, these populations are predicted to evolve to increase growth at the expense of defense. B) Least squares means +/- s.e.m. observed for native and invading genotypes of yellow starthistle for immune function (peroxidase activity in the presence of bacteria) and for seedling size. C) Least squares means for immune function and plant size in native populations. Solid lines show significant effects of size and bacterial strain. D) Least squares means for immune function and plant size in invading populations. Dashed lines show nonsignificant effects of size and significant effects of bacterial strain. All absorbance values for peroxidase activity were measured in this study. Absorbance values are based upon the five bacterial strains eliciting significant regional differences in peroxidase activity. All size measurements are for plant genotypes from same populations used in this study, measured in a previous glasshouse study (Dlugosch et al. 2015).

## DISCUSSION

Using a recently developed high throughput assay of immune function, our study demonstrated consistent differences in the influence of both plant genotype and bacterial strain on plant-pathogen interactions. We found that yellow starthistle genotypes from the native European range have a higher immune response to bacterial infection than genotypes found in the invaded California (USA) range. These patterns were consistent with a potential negative trade-off between immune response and growth in this highly invasive species.

Our study focused on measuring the oxidative burst as a result of PTI. Hormonal crosstalk between PTI and growth may explain trade-offs between defense and growth in plants. Three hormones, auxins, brassinosteroids, and gibberellins may play an important role in this crosstalk (Huot et al., 2014). Auxins regulate plant growth, stem and petiole elongation and root structure growth (Santner and Estelle, 2009; Vanneste and Friml, 2009; Kieffer et al., 2010; Kazan, 2013). Pathogens can synthesize their own auxin or alter the plant’s auxin synthesis to promote infection (Yamada, 1993; Glickmann et al., 1998; Abramovitch et al., 2003; O’Donnell et al., 2003; Kidd et al., 2011). When PTI is triggered the plant will actively suppress auxin signaling, inhibiting plant growth (Navarro et al., 2004). Brassinosteroids influence growth at every stage of a plant’s development (Gruszka, 2013; Hao et al., 2013; Fariduddin et al., 2014) and gibberellins influence growth, flower initiation and development (Sun, 2011). When brassinosteroids and gibberellins have elevated signaling, PTI is suppressed (Albrecht et al., 2012; Belkhadir et al., 2012). These hormonal functions create direct trade-offs between defense and growth functions, such that changes in a plant’s environment may favor one function over the other.

We found evidence supporting differential immune response to bacterial strains between native and invaded ranges in yellow starthistle. For all bacterial strains that showed a significant difference between regions, native range genotypes had higher immune response than the invaded range genotypes. Peroxidase activity was particularly high among genotypes from western Europe, which is the source of the invading populations that we studied here (Barker et al., 2017), indicating that invaders are particularly divergent from their progenitors. Coupled with the observation that invading genotypes from these populations also grow larger under a common environment (Dlugosch et al., 2015), our data support the hypothesis of a growth-defence trade-off at the regional scale. We also found that variation in immune response within the native range was significantly negatively related to plant size at the population scale, which is consistent with a trade-off and further suggests that there is standing variation along this axis in the native range. These results support the idea that introduced species may evolve along this type of trade-off, potentially increasing their invasiveness as a result of reduced investment in defenses (Blossey and Notzold, 1995; Maron et al., 2004; Bossdorf et al., 2005; Blumenthal and Hufbauer, 2007; Blumenthal et al., 2009; Felker-Quinn et al., 2013). To ascertain whether this has indeed been the case for yellow starthistle, further studies are needed to understand the prevalence of disease in different parts of its range, and their effects on fitness. Even if yellow starthistle has not escaped pathogens in this invasion, invaders may still evolve increased fitness at the expense of defense if they can better tolerate attack, as might be the case if they have access to more resources (Blumenthal, 2006; Chun et al., 2010).

In contrast to the pattern that we observed among native genotypes, there was no apparent relationship between plant size and immune responses among invading genotypes. Instead, immune response appeared to remain at a low level across a wide variation in plant size. This pattern could reflect a minimal wounding and/or pathogen response for the species. Wounding alone induces a response that strongly resembles pathogen attack and will stimulate ROS production similar to PTI (Baron and Zambryski, 1995). Regardless, our data suggest that some increases in plant size seen among invaders do not come at additional cost of the peroxidase response in immune function.

These plant-pathogen interactions included consistent differences in the immune response to the different bacterial strains used in our experiments, including particularly strong reactions to known plant pathogens. The highest peroxidase activities were observed in response to two strains of *Curtobacterium flaccumfaciens* (1038,1043). *Curtobacterium flaccumfaciens* is a known plant pathogen responsible for bacterial wilt (Hedges, 1922; González et al., 2005). The next highest levels of activity were seen in response to *Pseudomonas asturiensis* (1033) and *P. viridiflava* (1031). *Pseudomonas asturiensis* appears to be the cause of a newly recognized disease on soybean leaves (González et al., 2012, 2013). *Pseudomonas viridiflava* is a plant pathogen commonly infecting *Arabidopsis thaliana* and many agricultural crops (Billing, 1970; Wilkie et al., 1973; Goumans and Chatzaki, 1998; Jakob et al., 2002; Goss and Bergelson, 2007). The differential immune response among strains suggests different levels of pathogenicity, potentially due to variation in immune suppression (Abramovitch and Martin, 2004). Alternatively, live strains may have varied in their rate of growth during incubation, resulting in different levels of exposure to plant cells. Whole-plant inoculation experiments, allowing tests of bacterial growth within the plant and their putative fitness, would be valuable for assessing pathogenicity differences both among strains and between infections of invading and native genotypes.

In general, plants that have encountered particular bacterial pathogens in their evolutionary history might be especially well-adapted to recognize and respond to these strains, though the bacteria may evolve their own abilities to suppress locally-adapted immune responses (Karasov et al., 2014/4; Thompson and Burdon, 1992; Gilbert, 2002; Gilbert and Parker, 2010). We selected pathogens from the native range to assay potential escape from these natural enemies, but they may also reflect more recent evolution of local interactions. Our strains were from western European populations, and there were two cases of matches of plants and bacteria from the same collection sites. In general, western European genotypes were most responsive in the peroxidase assay, consistent with a locally-adapted response for the plants. In the case of the strain collected from site ‘CAZ’ (1032), genotypes originating from this site ranked the second highest in their response to this local strain, but this was a typical ranking for these plant genotypes across all strains. In the case of the strain collected from site ‘SAZ’ (1038), plant genotypes from this site demonstrated one of their highest responses relative to other populations (though differences among native plant populations were not significant).

Our invading plant genotypes also showed significant differences among source populations. Genotypes from the coastal area (DIA, GIL), closest to the site of introduction in the San Francisco Bay area, as well as a population near the eastern leading edge of the invasion (SIE) showed the lowest immune responses, while sites in the Central Valley of California tended to be more responsive (particularly RB). Whether these differences reflect variation in patterns of selection on defense traits is unknown. The invasion of yellow starthistle into the Central Valley is particularly severe (Pitcairn et al., 1997; Swope and Parker, 2010), and it will be important to understand whether these areas are experiencing higher pathogen incidence and/or whether immune responses and growth might be benefitting simultaneously from adaptations to higher resource available in these areas (Dlugosch et al., 2015).

Our experiments revealed several methodological considerations worth noting. Bacterial elicitors should be present in non-living preparations of bacterial cultures or elicitor isolates, and such non-living material is often used as an efficient means of studying plant-bacteria interactions (Felix et al., 1999; Daudi et al., 2012; Mott et al., 2016). During our Experiment 1, however, differences were observed between the plants’ immune response to autoclaved strains and live strains. While the autoclaved strains did elicit a mean increase in activity, this was not statistically different from leaf only controls without bacteria added. A live culture of one bacterial strain elicited a much stronger response in our experiment. Autoclaving may have denatured the elicitors to the extent that the plants could not recognize the PAMPs, or elicitor production may have been higher in bacteria incubated with leaf disks. Mechanically lysed bacteria, in place of autoclaved material, could be used to further clarify the impact of autoclaving on bacterial elicitors in this assay.

A particularly important methodological issue arose in our ‘Leaf Only’ treatments of leaf disks in sterile fresh media. This treatment showed elevated mean peroxidase activity over controls with no leaf disks, in both experiments, however in Experiment 2 this activity was also significantly higher than activity in all leaf treatments exposed to live bacterial strains. Some peroxidase activity is expected in the absence of bacteria due to responses to wounding (Minibayeva et al., 2015), but this would not explain a higher level of activity than leaf treatments that included bacteria. Peroxidase activity could have been additionally elevated due to incidental infection in our plants under glasshouse conditions before the plant tissue was harvested (Jones and Dangl, 2006), although any unintended exposure to pathogens should have again affected all leaf treatments equally (given that disks for controls and bacterial treatments were sampled from the same leaf at the same time).

We propose two potential explanations for these patterns. First, immune suppression by our bacterial treatments is possible. Plant pathogens have a variety of ways to suppress the immune response, including interfering with PTI and the hypersensitive response (Bretz et al., 2003; Abramovitch and Martin, 2004), and inhibiting the host’s programmed cell death (Abramovitch et al., 2003; Espinosa et al., 2003; Abramovitch and Martin, 2004). Pathogens can release effectors that interfere with PTI and some effectors can specifically target and inhibit peroxidase activity, as seen in corn smut (Hemetsberger et al., 2012) and infections of potatoes (Doke, 1975). It is possible that our experimental strains, derived from the native range of yellow starthistle, are capable of a greater degree of immune suppression in yellow starthistle than was a pathogen acquired from our glasshouse environment in the invaded range. In this case, our results would indicate that immune suppression is higher or host response weaker in our invading plant genotypes, both of which would indicate loss of defense function relative to native genotypes. Second, exposure of leaf disks to fresh media could elicit different peroxidase activity than the media remaining in active live cultures, such that fresh media is not an appropriate control in our experiment. Previous tests of peroxidase activity have used bacterial elicitors suspended in water, with water as the control treatment (Reuveni et al., 1992; Nawar and Kuti, 2003; Mott et al., 2016). If media influences the background wounding response, then the growth of bacteria in their own media could alter this response. In the immediate future, we are planning an experiment to incorporate treatments that filter bacteria from their media, allowing us to test responses both media after bacterial growth, and to bacteria removed from any media. We note that differences in relative performance of live strain treatments and the ‘Leaf Only’ treatment in our two experiments indicates that either this response to media or immune suppression is not acting equally across experiments in our system.

Our experiments did indicate consistent differences in immune activity between native and invading genotypes, providing valuable information for further studies of plant-pathogen interactions and potential growth-defense trade-offs in this system. Detailed studies of pathogen prevalence in the field and whole-plant inoculation experiments (that quantify fitness for both plant and pathogen) will be necessary to definitely identify whether introduced plants are benefitting from evolution along growth-defense trade-offs and whether this is the result of escape from native pathogens. We found that the high-throughput peroxidase assay provided an efficient method for exploring whether these intensive avenues of future research are warranted, and that a unique strength of this approach is its ability to replicate tests of plant-pathogen genotype interactions at the scale of a single leaf. This opportunity for replication is particularly important for non-clonal, obligate outcrossing species such as yellow starthistle, in which the same plant genotype cannot be replicated at the whole-plant scale. In our case, we were able to identify signatures of shifts in plant functions that are hypothesized to be integral to colonization success and the evolution of highly invasive plant species.

## ACKNOWLEDGEMENTS

We thank J. Aspinwall and A. Davis for help with peroxidase assays, J. Braasch and M. Johnson for help with planting, and A. Mott for help with protocol development. This study was supported by USDA grant 2015-67013-23000 to KMD and DAB.

## LITERATURE CITED

Abramovitch, R.B., Y.-J. Kim, S. Chen, M.B. Dickman, and G.B. Martin. 2003. Pseudomonas type III effector AvrPtoB induces plant disease susceptibility by inhibition of host programmed cell death. The EMBO journal 22: 60–69.

Abramovitch, R.B., and G.B. Martin. 2004. Strategies used by bacterial pathogens to suppress plant defenses. Current Opinion in Plant Biology 7: 356–364.

Albrecht, C., F. Boutrot, C. Segonzac, B. Schwessinger, S. Gimenez-Ibanez, D. Chinchilla, J.P. Rathjen, et al. 2012. Brassinosteroids inhibit pathogen-associated molecular pattern-triggered immune signaling independent of the receptor kinase BAK1. Proceedings of the National Academy of Sciences, USA 109: 303–308.

Andonian, K., and J.L. Hierro. 2011. Species interactions contribute to the success of a global plant invader. Biological Invasions 13: 2957–2965.

Andonian, K., J.L. Hierro, L. Khetsuriani, P.I. Becerra, G. Janoyan, D. Villareal, L.A. Cavieres, et al. 2012. Geographic mosaics of plant-soil microbe interactions in a global plant invasion. Journal of Biogeography 39: 600–608.

Andonian, K., J.L. Hierro, L. Khetsuriani, P. Becerra, G. Janoyan, D. Villarreal, L. Cavieres, et al. 2011. Range-Expanding Populations of a Globally Introduced Weed Experience Negative Plant-Soil Feedbacks. PloS One 6: e20117.

Barker, B.S., K. Andonian, S.M. Swope, D.G. Luster, and K.M. Dlugosch. 2017. Population genomic analyses reveal a history of range expansion and trait evolution across the native and invaded range of yellow starthistle (*Centaurea solstitialis*). Molecular Ecology 26: 1131–1147.

Baron, C., and P.C. Zambryski. 1995. The plant response in pathogenesis, symbiosis, and wounding: variations on a common theme? Annual review of genetics 29: 107–129.

Belkhadir, Y., Y. Jaillais, P. Epple, E. Balsemão-Pires, J.L. Dangl, and J. Chory. 2012. Brassinosteroids modulate the efficiency of plant immune responses to microbe-associated molecular patterns. Proceedings of the National Academy of Sciences, <u>USA</u> 109: 297–302.

Billing, E. 1970. *Pseudomonas viridiflava* (Burkholder, 1930; Clara 1934). Journal of Applied Bacteriology 33: 492–500.

Blossey, B., and R. Notzold. 1995. Evolution of Increased Competitive Ability in Invasive Nonindigenous Plants: A Hypothesis. Journal of Ecology 83: 887.

Blumenthal, D.M. 2006. Interactions between resource availability and enemy release in plant invasion. Ecology Letters 9: 887–895.

Blumenthal, D.M., and R.A. Hufbauer. 2007. Increased plant size in exotic populations: a common-garden test with 14 invasive species. Ecology 88: 2758–2765.

Blumenthal, D., C.E. Mitchell, P. Pysek, and V. Jarosík. 2009. Synergy between pathogen release and resource availability in plant invasion. Proceedings of the National Academy of Sciences, USA 106: 7899–7904.

Bossdorf, O., H. Auge, L. Lafuma, W.E. Rogers, E. Siemann, and D. Prati. 2005. Phenotypic and genetic differentiation between native and introduced plant populations. Oecologia 144: 1–11.

Bretz, J.R., N.M. Mock, J.C. Charity, S. Zeyad, C.J. Baker, and S.W. Hutcheson. 2003. A translocated protein tyrosine phosphatase of Pseudomonas syringae pv. tomato DC3000 modulates plant defence response to infection. Molecular Microbiology 49: 389–400.

Camejo, D., Á. Guzmán-Cedeño, and A. Moreno. 2016. Reactive oxygen species, essential molecules, during plant–pathogen interactions. Plant Physiology and Biochemistry 103: 10–23.

Chakravarthy, S., A.C. Velásquez, and G.B. Martin. 2009. Assay for Pathogen-Associated Molecular Pattern (PAMP)-Triggered Immunity (PTI) in Plants. Journal of visualized experiments: JoVE.

Chisholm, S.T., G. Coaker, B. Day, and B.J. Staskawicz. 2006. Host-microbe interactions: shaping the evolution of the plant immune response. Cell 124: 803–814.

Chun, Y.J., M. van Kleunen, and W. Dawson. 2010. The role of enemy release, tolerance and resistance in plant invasions: linking damage to performance. Ecology Letters 13: 937–946.

Colautti, R.I., A. Ricciardi, I.A. Grigorovich, and H.J. MacIsaac. 2004. Is invasion success explained by the enemy release hypothesis? Ecology Letters 7: 721–733.

Daudi, A., Z. Cheng, J.A. O’Brien, N. Mammarella, S. Khan, F.M. Ausubel, and G.P. Bolwell. 2012. The Apoplastic Oxidative Burst Peroxidase in Arabidopsis Is a Major Component of Pattern-Triggered Immunity. Plant Cell 24: 275–287.

Ditomaso, J.M., G.B. Kyser, and M.J. Pitcairn. 2006. Yellow Starthistle Management Guide.

Dlugosch, K.M., F. Alice Cang, B.S. Barker, K. Andonian, S.M. Swope, and L.H. Rieseberg. 2015. Evolution of invasiveness through increased resource use in a vacant niche. Nature Plants 1: 15066.

Dodds, P.N., and J. P. Rathjen. 2010. Plant immunity: towards an integrated view of plant-pathogen interactions. Nature Reviews Genetics 11: 539–548.

Doke, N. 1975. Prevention of the hypersensitive reaction of potato cells to infection with an incompatible race of Phytophthora infestans by constituents of the zoospores. Physiological Plant Pathology 7: 1–7.

Eriksen, R.L., T. Desronvil, J.L. Hierro, and R. Kesseli. 2012. Morphological differentiation in a common garden experiment among native and non-native specimens of the invasive weed yellow starthistle (*Centaurea solstitialis*). Biological Invasions 14: 1459–1467.

Espinosa, A., M. Guo, V.C. Tam, Z.Q. Fu, and J.R. Alfano. 2003. The Pseudomonas syringae type III-secreted protein HopPtoD2 possesses protein tyrosine phosphatase activity and suppresses programmed cell death in plants. Molecular Microbiology 49: 377–387.

Faillace, C.A., N.S. Lorusso, and S. Duffy. 2017. Overlooking the smallest matter: viruses impact biological invasions. Ecology Letters 20: 524–538

Fariduddin, Q., M. Yusuf, I. Ahmad, and A. Ahmad. 2014. Brassinosteroids and their role in response of plants to abiotic stresses. Biologia Plantarum 58: 9–17.

Felix, G., J.D. Duran, S. Volko, and T. Boller. 1999. Plants have a sensitive perception system for the most conserved domain of bacterial flagellin. Plant journal 18: 265–276.

Felker-Quinn, E., J.A. Schweitzer, and J.K. Bailey. 2013. Meta-analysis reveals evolution in invasive plant species but little support for Evolution of Increased Competitive Ability (EICA). Ecology and Evolution 3: 739–751.

Gerlach, J.D. 1997. How the West Was Lost: Reconstructing the Invasion Dynamics of Yellow Starthistle and Other Plant Invaders of Western Rangelands and Natural Areas. California Exotic Pest Plant Council Symposium Proceedings.

Gilbert, G.S. 2002. Evolutionary ecology of plant diseases in natural ecosystems. Annual Review of Phytopathology 40: 13–43.

Gilbert, G.S., and I.M. Parker. 2010. Rapid evolution in a plant-pathogen interaction and the consequences for introduced host species. Evolutionary Applications 3: 144–156.

Glickmann, E., L. Gardan, S. Jacquet, S. Hussain, M. Elasri, A. Petit, and Y. Dessaux. 1998. Auxin Production Is a Common Feature of Most Pathovars of Pseudomonas syringae. Molecular Plant-microbe Interactions 11: 156–162.

González, A.J., I. Cleenwerck, P. De Vos, and A.M. Fernández-Sanz. 2013. Pseudomonas asturiensis sp. nov., isolated from soybean and weeds. Systematic and Applied Microbiology 36: 320–324.

González, A.J., A.M. Fernández, M. San José, G. González-Varela, and M.R. Rodicio. 2012. A Pseudomonas viridiflava-related bacterium causes a dark-reddish spot disease in Glycine max. Applied and Environmental Microbiology 78: 3756–3758.

González, A.J., J.C. Tello, and M.R. Rodicio. 2005. Bacterial Wilt of Beans (*Phaseolus vulgaris*) Caused by *Curtobacterium flaccumfaciens* in Southeastern Spain. Plant Disease 89: 1361–1361.

Goss, E.M., and J. Bergelson. 2007. Fitness consequences of infection of Arabidopsis thaliana with its natural bacterial pathogen Pseudomonas viridiflava. Oecologia 152: 71–81.

Goumans, D.E., and A. K. Chatzaki. 1998. Characterization and host range evaluation of Pseudomonas viridiflava from melon, blite, tomato, chrysanthemum and eggplant. European Journal of Plant Pathology 104: 181–188.

Gruszka, D. 2013. The brassinosteroid signaling pathway-new key players and interconnections with other signaling networks crucial for plant development and stress tolerance. International Journal of Molecular Sciences 14: 8740–8774.

Hao, J., Y. Yin, and S.-Z. Fei. 2013. Brassinosteroid signaling network: implications on yield and stress tolerance. Plant Cell Reports 32: 1017–1030.

Hedges, F. 1922. A bacterial wilt of the Bean caused by Bacterium flaccumfaciens nov. sp. Science (Washington) 55: 433–434.

Hemetsberger, C., C. Herrberger, B. Zechmann, M. Hillmer, and G. Doehlemann. 2012. The Ustilago maydis effector Pep1 suppresses plant immunity by inhibition of host peroxidase activity. PLoS Pathogens 8: e1002684.

Huot, B., J. Yao, B.L. Montgomery, and S.Y. He. 2014. Growth–Defense Tradeoffs in Plants: A Balancing Act to Optimize Fitness. Molecular Plant 7: 1267–1287.

Jakob, K., E.M. Goss, H. Araki, T. Van, M. Kreitman, and J. Bergelson. 2002. Pseudomonas viridiflava and *P. syringae*—Natural Pathogens of *Arabidopsis thaliana*. Molecular Plant-microbe Interactions 15: 1195–1203.

Jones, J.D.G., and J.L. Dangl. 2006. The plant immune system. Nature 444: 323–329.

Karasov, T.L., M.W. Horton, and J. Bergelson. 2014/4. Genomic variability as a driver of plant–pathogen coevolution? Current Opinion in Plant Biology 18: 24–30.

Kazan, K. 2013. Auxin and the integration of environmental signals into plant root development. Annals of Botany 112: 1655–1665.

Keane, R.M., and M.J. Crawley. 2002. Exotic plant invasions and the enemy release hypothesis. Trends in Ecology & Evolution 17: 164–170.

Kidd, B.N., N.Y. Kadoo, B. Dombrecht, M. Tekeoglu, D.M. Gardiner, L.F. Thatcher, E.A.B. Aitken, et al. 2011. Auxin signaling and transport promote susceptibility to the root-infecting fungal pathogen Fusarium oxysporum in Arabidopsis. Molecular Plant-microbe Interactions 24: 733–748.

Kieffer, M., J. Neve, and S. Kepinski. 2010. Defining auxin response contexts in plant development. Current Opinion in Plant Biology 13: 12–20.

Kulmatiski, A., K.H. Beard, J.R. Stevens, and S.M. Cobbold. 2008. Plant–soil feedbacks: a meta-analytical review. Ecology Letters 11: 980–992.

Lamb, C., and R.A. Dixon. 1997. The Oxidative Burst in Plant Disease Resistance. Annual Review of Plant Physiology and Plant Molecular Biology 48: 251–275.

Lu-Irving, P., J. Harencar, H. Sounart, S.R. Welles, S.M. Swope, D.A. Baltrus, and K.M. Dlugosch. 2017. Escape from bacterial diversity: potential enemy release in invading yellow starthistle (Centaurea solstitialis) microbiomes. bioRxiv119917.

Maron, J.L., M. Vilà, and J. Arnason. 2004. Loss of enemy resistance among introduced populations of St. John’s Wort (Hypericum perforatum). Ecology 85: 3243–3253.

Medzhitov, R. 2013. Pattern recognition theory and the launch of modern innate immunity. Journal of Immunology 191: 4473–4474.

Minibayeva, F., R.P. Beckett, and I. Kranner. 2015. Roles of apoplastic peroxidases in plant response to wounding. Phytochemistry 112: 122–129.

Mitchell, C.E., D. Blumenthal, V. Jarošík, E.E. Puckett, and P. Pyšek. 2010. Controls on pathogen species richness in plants’ introduced and native ranges: roles of residence time, range size and host traits. Ecology Letters 13: 1525–1535.

Mitchell, C.E., and A.G. Power. 2003. Release of invasive plants from fungal and viral pathogens. Nature 421: 625–627.

Mott, G.A., S. Thakur, E. Smakowska, P.W. Wang, Y. Belkhadir, D. Desveaux, D.S. Guttman, et al. 2016. Genomic screens identify a new phytobacterial microbe-associated molecular pattern and the cognate Arabidopsis receptor-like kinase that mediates its immune elicitation. Genome Biology 17: 98.

Navarro, L., C. Zipfel, O. Rowland, I. Keller, S. Robatzek, T. Boller, and J.D.G. Jones. 2004. The transcriptional innate immune response to flg22. Interplay and overlap with Avr gene-dependent defense responses and bacterial pathogenesis. Plant Physiology 135: 1113–1128.

Nawar, H.F., and J.O. Kuti. 2003. Wyerone Acid Phytoalexin Synthesis and Peroxidase Activity as Markers for Resistance of Broad Beans to Chocolate Spot Disease. Journal of Phytopathology 151: 564–570.

O’Donnell, P.J., E.A. Schmelz, P. Moussatche, S.T. Lund, J.B. Jones, and H.J. Klee. 2003. Susceptible to intolerance--a range of hormonal actions in a susceptible Arabidopsis pathogen response. Plant Journal 33: 245–257.

Pejchar, L., and H.A. Mooney. 2009. Invasive species, ecosystem services and human well-being. Trends in Ecology & Evolution 24: 497–504.

Pimentel, D., R. Zuniga, and D. Morrison. 2005. Update on the environmental and economic costs associated with alien-invasive species in the United States. Ecological Economics 52: 273–288.

Pitcairn, M.J., R.A. O’Connell, and J.M. Gendron. 1997. Yellow starthistle: survey of statewide distribution. Biological control program annual summary 64–66.

Pitcairn, M.J., S. Schoenig, R. Yacoub, and J. Gendron. 2006. Yellow starthistle continues its spread in California. California Agriculture 60: 83–90.

Reuveni, R., M. Shimoni, Z. Karchi, and J. Kuć. 1992. Peroxidase activity as a biochemical marker for resistance of Muskmelon (Cucumis melo) to Pseudoperonospora cubensis. Phytopathology 82: 749–753.

Santner, A., and M. Estelle. 2009. Recent advances and emerging trends in plant hormone signalling. Nature 459: 1071–1078.

Sun, T.-P. 2011. The molecular mechanism and evolution of the GA-GID1-DELLA signaling module in plants. Current Biology 21: R338–45.

Swope, S.M., and I.M. Parker. 2010. Widespread seed limitation affects plant density but not population trajectory in the invasive plant Centaurea solstitialis. Oecologia 164: 117–128.

Thompson, J.N., and J.J. Burdon. 1992. Gene-for-gene coevolution between plants and parasites. Nature 360: 121–125.

Vanneste, S., and J. Friml. 2009. Auxin: a trigger for change in plant development. Cell 136: 1005–1016.

Widmer, T.L., F. Guermache, M.Y. Dolgovskaia, and S.Y. Reznik. 2007. Enhanced Growth and Seed Properties in Introduced vs. Native Populations of Yellow Starthistle (Centaurea solstitialis). Weed Science 55: 465–473.

Wilkie, J.P., D.W. Dye, and D.R.W. Watson. 1973. Further hosts of *Pseudomonas viridiflava*. New Zealand Journal of Agricultural Research 16: 315–323.

Win, J., A. Chaparro-Garcia, K. Belhaj, D.G.O. Saunders, K. Yoshida, S. Dong, S. Schornack, et al. 2012. Effector biology of plant-associated organisms: concepts and perspectives. Cold Spring Harbor symposia on quantitative biology 77: 235–247.

Wojtaszek, P. 1997. Oxidative burst: an early plant response to pathogen infection. The Biochemical Journal 322: 681–692.

Yamada, T. 1993. The role of auxin in plant-disease development. Annual Review of Phytopathology 31: 253–273.

